# Evolutionarily conserved regulation of embryonic fast-twitch skeletal muscle differentiation by Pbx factors

**DOI:** 10.1101/2020.02.21.960484

**Authors:** Gist H. Farr, Bingsi Li, Maurizio Risolino, Nathan M. Johnson, Zizhen Yao, Robert M. Kao, Mark W. Majesky, Stephen J. Tapscott, Licia Selleri, Lisa Maves

**Author notes:** Corresponding author: Lisa Maves, Center for Developmental Biology and Regenerative Medicine, Seattle Children’s Research Institute, 1900 Ninth Avenue, Seattle, WA 98101 USA.

## Abstract

Vertebrate skeletal muscles are composed of both slow-twitch and fast-twitch fiber types. How the differentiation of distinct fiber types is activated during embryogenesis is not well characterized. Skeletal muscle differentiation is initiated by the activity of the myogenic basic helix-loop-helix (bHLH) transcription factors Myf5, Myod1, Myf6, and Myog. Myod1 functions as a muscle master regulatory factor and directly activates muscle differentiation genes, including those specific to both slow and fast muscle fibers. Our previous studies showed that Pbx TALE-class homeodomain proteins bind with Myod1 on the promoter of the zebrafish fast muscle gene *mylpfa* and are required for proper activation of *mylpfa* expression and the fast-twitch muscle-specific differentiation program in zebrafish embryos. Pbx proteins have also been shown to bind regulatory regions of muscle differentiation genes in mammalian muscle cells in culture. Here, we use new zebrafish mutant strains to confirm the essential roles of zebrafish Pbx factors in embryonic fast muscle differentiation. Furthermore, we examine the requirements for *Pbx* genes in mouse embryonic skeletal muscle differentiation, an area that has not been investigated in the mammalian embryo. Removing *Pbx1* function from skeletal muscle in *Myf5^Cre/+;^Pbx1^fl/fl^* mouse embryos has minor effects on embryonic muscle development. However, concomitantly deleting *Pbx2* function in *Myf5^Cre/+^;Pbx1^fl/fl^;Pbx2^-/-^* mouse embryos causes delayed activation and reduced expression of fast muscle differentiation genes. In the mouse, *Pbx1/Pbx2*-dependent fast muscle genes closely match those that have been previously shown to be dependent on murine *Six1* and *Six4*. This work establishes evolutionarily conserved requirements for Pbx factors in embryonic fast muscle differentiation. Our studies are revealing how Pbx homeodomain proteins help direct specific cellular differentiation pathways.

## Introduction

Skeletal muscle fiber diversity allows for a range of movements, activity and metabolism, but muscle fiber diversity can also affect susceptibility to muscle disease and muscle wasting (reviewed in Schiaffino and Reggiani, 2011; and Talbot and Maves, 2016). There are two main types of skeletal muscle fibers: slow twitch and fast twitch. Understanding how skeletal muscle fiber type differentiation is controlled is important for addressing skeletal muscle defects that occur in muscle aging, metabolic diseases, and muscular dystrophies (reviewed in Ciciliot et al., 2013; Ljubicic et al., 2014; and Talbot and Maves, 2016). Many processes have been described that regulate the plasticity of skeletal muscle fiber-type identities during post-natal periods (reviewed in Bassel-Duby and Olson, 2006; Blaauw et al., 2013; and Schiaffino and Reggiani, 2011). However, the mechanisms that establish muscle fiber-type identities during embryogenesis are less well understood.

Skeletal muscle differentiation is initiated in embryogenesis by the myogenic regulatory factor (MRF) family of basic helix-loop-helix (bHLH) transcription factors: Myf5, Myod1, Myf6, and Myog (reviewed in Hernández-Hernández et al, 2017; Lassar, 2017; and Zammit, 2017). The activity of these factors is highly conserved. In both mice and zebrafish, the MRFs are required for skeletal muscle myogenesis (reviewed in Buckingham and Vincent, 2009; and Rossi and Messina, 2014). In particular, Myod1 functions as a muscle master regulatory factor and controls skeletal muscle differentiation through DNA binding and transcriptional activation of broad gene expression programs, including genes encoding skeletal muscle contractile proteins (Blais et al., 2005; Cao et al., 2006; Cao et al., 2010; Pliner et al., 2018; Soleimani et al., 2012). In the differentiation of specific muscle fiber types, Myod1 has been associated with promoting fast muscle differentiation. In mouse muscle, Myod1 is expressed more strongly in fast fibers than in slow fibers, and gain- and loss-of-function experiments in mice and zebrafish have shown that Myod1 promotes fast fiber types (Ekmark et al., 2007; Hammond et al., 2007; Hinits et al., 2011; Hughes et al., 1997; Macharia et al., 2010; Maves et al., 2007; Seward et al., 2001). However, these and other studies have found that Myod1 is also needed for slow fiber differentiation and directly activates both slow and fast muscle differentiation genes (Blais et al., 2005; Cao et al., 2006; Cao et al., 2010; Hinits et al., 2011; Hughes et al., 1997; Maves et al., 2007; Seward et al., 2001; Soleimani et al., 2012). Myod1’s DNA binding and transcriptional activity can be modulated, positively and negatively, by many other transcription factors (reviewed in Berkes and Tapscott, 2005; Fong and Tapscott, 2013; and Talbot and Maves, 2016). Understanding how the activity of MRFs such as Myod1 is regulated during embryogenesis should provide insight into skeletal muscle fiber type specification (Talbot and Maves, 2016).

We already understand some components of a transcriptional network used to activate skeletal muscle fiber-type specification during embryogenesis (reviewed in Jackson and Ingham, 2013; and Rossi and Messina, 2014). Transcription factors of the Six homeodomain family are needed to promote MRF gene expression and muscle differentiation in both mice and zebrafish embryos (Bessarab et al., 2008; Grifone et al., 2005; Niro et al., 2010; Nord et al., 2013; O’Brien et al., 2014; Richard et al., 2011; Talbot et al., 2019). In mouse embryos with compound loss of *Six1;Six4*, and in zebrafish embryos with morpholino knock-down of *six1a* and *six1b*, fast muscle gene expression is reduced in the early myotome, whereas expression of slow muscle genes is largely not affected (Bessarab et al., 2008; Niro et al., 2010; O’Brien et al., 2014). However, in zebrafish embryos with genetic deletion of all four zebrafish *six1* and *six4* genes, migratory muscle precursor cells do not undergo proper specification or migration, but embryonic fast muscle in the trunk still forms normally (Talbot et al., 2019). The Sox6 transcription factor promotes fast muscle differentiation by repressing slow muscle differentiation in zebrafish embryos and in mouse embryos (Hagiwara et al., 2005; Hagiwara et al., 2007; Jackson et al., 2015; von Hofsten et al., 2008). In zebrafish, the transcription factor Prdm1a acts as a critical switch in embryonic muscle fiber-type specification. Prdm1a directly represses fast muscle differentiation genes and also represses *sox6* expression, thereby promoting slow muscle differentiation in zebrafish (von Hofsten et al., 2008). However, even though *Prdm1* is expressed in the embryonic mouse myotome, the onset of myogenesis and fast and slow muscle differentiation appear to proceed normally in *Prdm1* mutant mouse embryos (Vincent et al., 2012). Thus, while there are many conserved aspects of muscle fiber-type specification and differentiation, there also appear to be differences between zebrafish and mouse in the regulation of embryonic muscle fiber type.

We previously showed, in zebrafish embryos, that Pbx TALE-class homeodomain proteins are required for the fast muscle differentiation program (Maves et al., 2007; Yao et al., 2013). In zebrafish embryos lacking Pbx proteins through antisense morpholino knockdown, activation of fast muscle differentiation genes, such as *mylpfa* and *atp2a1a*, is delayed and, even upon activation, these genes show reduced expression (Maves et al., 2007; Yao et al., 2013). Studies in zebrafish and mouse embryos, as well as in mammalian cell culture models, have shown that Pbx proteins bind with Myod1 on the promoters of a subset of Myod1 target genes, including *Myog* and *mylpfa* (Berkes et al., 2004; Cho et al., 2015; de la Serna et al., 2005; Dell’Orso et al., 2016; Heidt et al., 2007; Maves et al., 2007; Pliner et al., 2018). Pbx proteins can bind silent Myod1 target gene promoters prior to Myod1 binding and muscle differentiation, suggesting the potential for Pbx proteins to function as pioneer factors in skeletal muscle differentiation (Berkes et al., 2004; Cho et al., 2015; de la Serna et al., 2005; Dell’Orso et al., 2016; for discussion of Pbx proteins as pioneer factors, see Grebbin and Schulte, 2017 and Selleri et al., 2019). *Pbx* genes are broadly expressed during embryogenesis and are required for many aspects of mouse embryo development (reviewed in Moens and Selleri, 2006; and Selleri et al., 2019). However, these previous studies have not directly addressed the requirements for Pbx proteins in mammalian skeletal muscle development. Considering that the mechanisms underlying muscle fiber-type differentiation can vary between mice and zebrafish, we tested whether Pbx proteins are required for fast muscle differentiation during mouse embryogenesis.

Here we generate new zebrafish mutant strains to confirm that zebrafish embryos lacking Pbx expression show reduced and delayed activation of fast muscle differentiation genes. Furthermore, we investigate the requirements for Pbx factors in early mouse muscle development. We show that both Pbx1 and Pbx2 are broadly expressed in muscle precursors in embryonic mouse somites. We use a conditional knock-out approach to generate mouse embryos lacking both Pbx1 and Pbx2 in *Myf5*-expressing muscle precursors. We uncover that *Myf5^Cre/+^;Pbx1^fl/fl^;Pbx2^-/-^* embryos show reduced and delayed activation of fast muscle differentiation genes. Our findings demonstrate evolutionary conservation in the requirements for Pbx proteins in promoting activation of the fast muscle differentiation program during vertebrate embryogenesis.

## Materials and Methods

### Maintenance of animal strains and generation of zebrafish pbx2 mutant strain

All experiments involving live zebrafish and mice were carried out in compliance with Seattle Children’s Research Institute IACUC guidelines.

Zebrafish were raised and staged as previously described (Westerfield, 2000). Time (hpf) refers to hours post-fertilization at 28.5°C. In some cases, embryos were raised for periods at 24°C. For studies prior to 24 hpf, somite (s) number was used for staging, and mutant and control embryos were somite-stage matched. The wild-type stock and genetic background used was AB. The *pbx4^b557^* mutant strain was previously described and is likely a null allele (Pöpperl et al., 2000). *pbx4^b557^* genotyping was performed as previously described (Kao et al., 2015). The *pbx2^scm10^* mutant strain was generated with CRISPR/Cas9, using procedures we have previously described (Supplemental Figure 1; Farr et al., 2018). The *pbx2*-targeting single-guide construct was made by annealing the following oligonucleotides and ligating the resulting fragment into BsaI-digested pDR274 (Hwang et al., 2013): 5’ – [phos]TAGGTGTAGCCGGGCCAGAGAA – 3’ and 5’ – [phos]AAACTTCTCTGGCCCGGCTACA – 3’. To genotype the *pbx2^scm10^* strain, a dCAPS assay was designed (Neff et al., 1998) such that amplification of the mutant allele generated an XcmI site. The sequence of the *pbx2* CRISPR target site is shown in Supplemental Figure 1, along with the sequence of the *pbx2^scm10^* allele and its conceptual translation. Genotyping primers are provided in Table 1.

**Table 1.**
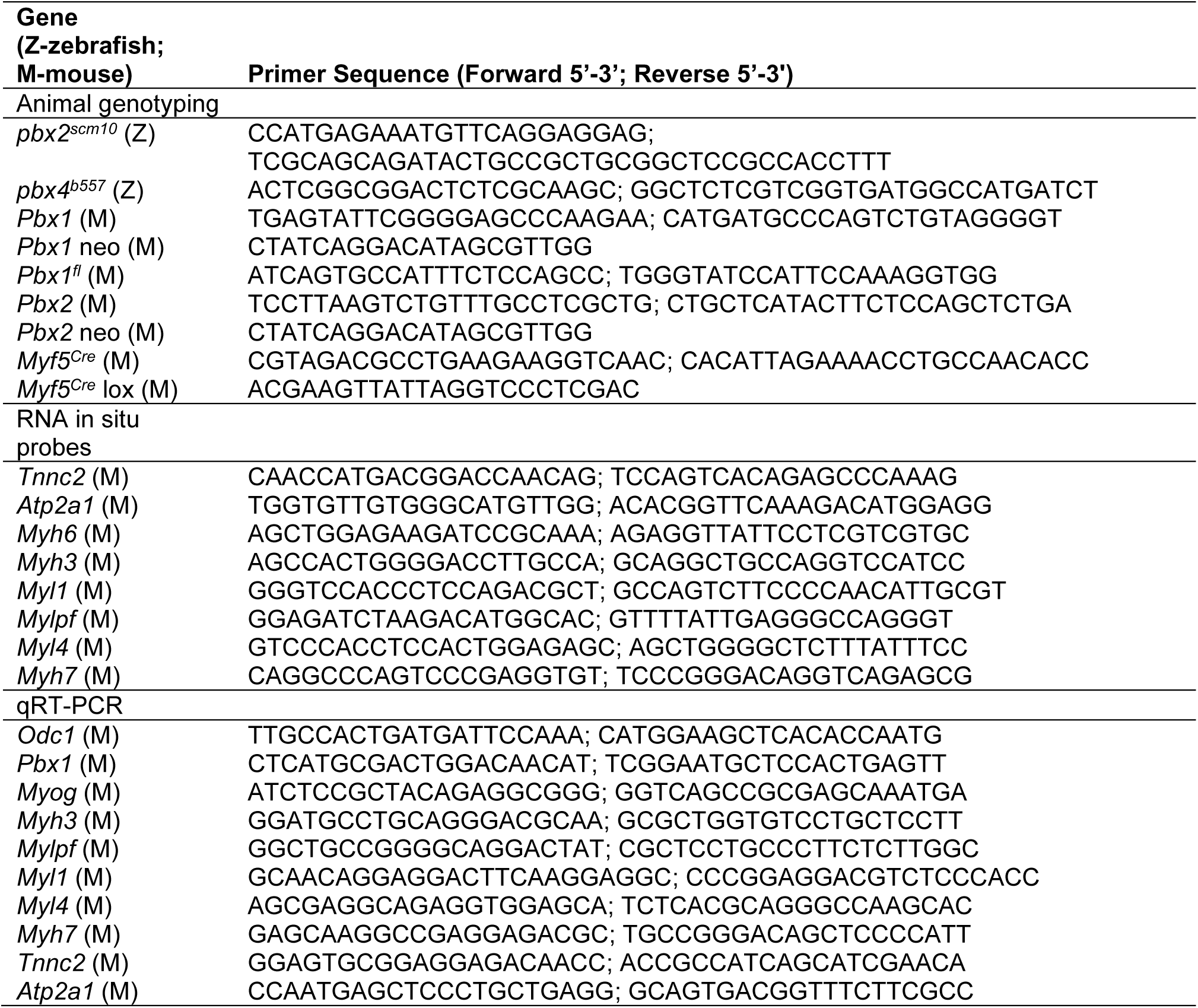
Primers used

The mouse strains used were previously described: the *Pbx1* knock-out allele (Selleri et al., 2001; hereafter called *Pbx1*^-/-^), the *Pbx1* conditional allele (*Pbx1^Δneo;Δex3^*; Koss et al., 2012; hereafter called *Pbx1^fl^*), the *Pbx2* knock-out allele (Selleri et al., 2004; hereafter called *Pbx2^-/-^*), and the *Myf5 Cre* allele (*Myf5^tm3(cre)Sor^*; Tallquist et al., 2000; hereafter called *Myf5^Cre^*; kindly provided by P. Soriano). Intercrosses were used to combine strains, which were maintained on a largely C57BL/6 background. Mouse embryos were obtained through timed matings, designating E0.5 as the morning on the day a vaginal plug was observed. Genotyping was performed by PCR using genomic DNA isolated from embryonic sacs or tail tissue. Protocols for *Pbx1*, *Pbx1^fl^*, *Pbx2*, and *Myf5^Cre^* genotyping were previously described (Koss et al., 2012; Selleri et al., 2001; Tallquist et al., 2000). Genotyping primers are provided in Table 1.

### Immunocytochemistry and RNA in situ hybridization

Zebrafish whole-mount RNA in situ hybridizations were performed on somite-stage-matched embryos as previously described (Maves et al., 2007; Talbot et al., 2010). The following cDNA probes were used: *krox-20* (*egr2b*-Zebrafish Information Network, Ruzicka et al., 2019; Oxtoby and Jowett, 1993); *myog* (Weinberg et al., 1996); *smyhc1* (Bryson-Richardson et al., 2005); *atp2a1* (Maves et al., 2007); *myl1* (Yao et al., 2013); *mylpfa* (formerly *mylz2*; Xu et al., 2000). Zebrafish embryos were imaged using a Leica TCS SP5 confocal microscope. Images were assembled using Adobe Photoshop.

For anti-Pbx1/Myf5 staining in mouse embryos, E11.5 embryos were fixed in 4% PFA/1X PBS overnight at 4^⁰^C and then dehydrated and stored in methanol at −20^⁰^C. Embryos were rehydrated using a methanol series performed at room temperature. Rehydrated embryos were washed in 1X PBST (1X PBS + 0.1% Triton X-100), followed by a puncture of the dorsal end of the neural tube to facilitate antibody diffusion. Embryos were rinsed in 1X PBSDT (1X PBS, 1% DMSO, 0.1% Triton X-100) followed by permeabilization in 1X PBS + 2% TritonX-100 for 30 minutes at room temperature. Permeabilized embryos were then rinsed in 1X PBSDT and blocked overnight in 10% normal donkey serum in 1X PBSDT. 1:100 rabbit polyclonal anti-Myf5 (IgG_H+L_, Santa Cruz, sc-302) and 1:100 monoclonal mouse anti-Pbx1b (IgG_1_, Santa Cruz, 41.1, Jacobs et al., 1999) were incubated in 2% normal donkey serum in 1X PBSDT for 2 days at 4^⁰^C. Embryos were then washed for five hours at room temperature in 1X PBSDT. For secondary antibodies, Alexa 488 donkey anti-mouse IgG_1_ and Alexa 594 donkey anti-rabbit IgG_H+L_ were used at 1:500 dilution in 2% normal donkey serum in 1X PBSDT overnight at 4^⁰^C. After secondary antibody staining, embryos were washed in 1X PBSDT. After post-fixation, embryos were pre-incubated in 30% glycerol + 0.1% (v/v) βScale2 (4M urea, 50% glycerol in distilled water) for 45 minutes; then *Clear^T2^* in 25% PEG/10% formamide was used for 1 hour at room temperature, followed by overnight in 50% PEG/20% formamide. Cleared embryos were mounted in 0.5% agarose gel in water and settled until gel polymerized; then *Clear^T2^* in 50% PEG/20% formamide was applied and the specimen was overlain with a coverslip. Optical stacks were collected using an upright Olympus single photon confocal microscope (20x water dipping objective, NA=0.95). Image datasets were analyzed using Fiji/ImageJ software.

For anti-Pbx2/Myod1 staining in mouse embryos, E11.5 embryos were fixed in pre-chilled 4% PFA/1X PBS for one hour. Fixed embryos were incubated in 30% sucrose/1X PBS and embedded in OCT frozen blocks. 30-micron-thick sagittal sections were taken on a Leica cryostat machine. Cryosections were blocked with 3% BSA/1% normal donkey serum in a humidified slide box and subsequently incubated with anti-Pbx2 (1:100, polyclonal rabbit, Santa Cruz, G-20) and anti-Myod1 (5.8A, mouse monoclonal IgG_1_) diluted in block solution. After rinses in 1X PBS with 0.25% Triton X-100, sections were then incubated with 1:500 Alexa 488 mouse IgG_1_ and Alexa 594 rabbit secondary antibodies in block solution for 1.5 hours at room temperature. Fluorescently labeled specimens were mounted in 2% n-propyl-gallate in 100% glycerol. Images were collected using a Leica SP5 inverted fluorescent confocal microscope with 20x air objective (NA=0.7) or 40x oil objective (NA=1.4). Fiji/ImageJ software was used for image analysis.

Mouse whole-mount RNA in situ hybridizations were performed on somite-stage-matched embryos as previously described (Di Giacomo et al., 2006). *Myog* probe was synthesized from a full-length mouse *Myog* cDNA cloned into the BamHI site of pCS2+. *Tnnc2* probe was synthesized from a 600bp cDNA fragment cloned into pCRII-TOPO. *Atp2a1* probe was synthesized from a 1236bp cDNA fragment cloned into pCRII-TOPO. *Myh6* probe was synthesized from a 1243bp cDNA fragment cloned into pCRII-TOPO. Additional cDNA probes for *Myh3*, *Myl1*, *Mylpf*, *Myl4*, and *Myh7* were generated by adding T7 promoters to PCR-amplified cDNA products. Primers used to amplify cDNA probes are provided in Table 1.

### Quantitative (q) RT-PCR

For mouse embryo qRT-PCR, total RNA samples were prepared from whole, individual, genotyped mouse embryos by disrupting the embryos in TRIzol (Ambion; Thermo Fisher Scientific) by trituration through a 16G needle followed by a 26G needle, after which the standard TRIzol protocol was followed. The isopropanol-precipitated RNA was solubilized in water and then further purified using an RNeasy Kit (Qiagen). 100 ng of RNA samples plus random hexamers were used in a reverse-transcriptase reaction with SuperScriptII Reverse Transcriptase (Invitrogen) or with the iScript cDNA Synthesis Kit (Bio-Rad). Real-time PCR was performed using SYBR Green (Bio-Rad) and an Applied Biosystems 7900HT System, or using the KAPA SYBR FAST kit (KAPA Biosystems) on a Bio-Rad CFX96 machine, according to the manufacturer’s instructions. Expression levels of the genes of interest were normalized to *Ornithine decarboxylase 1* (*Odc1*). For primers, see Table 1.

For each gene tested, a ΔΔC_t_ value was determined for each of the control/mutant pairs of sibling embryos. Two-tailed, unpaired Student’s t-tests with Welch’s correction were used to determine if the observed values differed significantly from the null hypothesis of ΔΔC_t_ values of 0. The resulting P values are shown in the graphs in Figures 2 and 4. To construct the graphs, the tables of ΔΔC_t_ values were log transformed to give fold differences (2^-ΔΔCt^). The boxes for each gene in the graphs extend from the 25th to 75th percentiles, the whiskers are at the minimum and maximum, the bar within the box represents the median, and the “+” symbol represents the mean. Statistical analysis was performed and the plots were made in GraphPad Prism7.

### RNA-seq library preparation and sequencing

Total RNA samples were obtained, using TRIzol Reagent (Life Technologies) and then cleaned using RNeasy Kit (Qiagen), from three *Myf5^Cre/+^;Pbx1^fl/fl^;Pbx2^-/-^* mutants and three *Pbx1^fl/+^;Pbx2^-/-^* control sibling whole mouse embryos at E11.5. Library preparation and sequencing was carried out by the FHCRC Genomics Shared Resource. Sequencing libraries were prepared from total RNA using the TruSeq RNA Sample Prep Kit (Illumina) according to the manufacturer’s instructions. Library size distributions were validated using an Agilent 2100 Bioanalyzer. Additional library quality control, blending of pooled indexed libraries, and cluster optimization were performed using the QPCR NGS Library Quantization Kit (Agilent Technologies). RNA-seq libraries were pooled (4-plex) and clustered onto a flow cell lane using an Illumina cBot. Sequencing occurred on an Illumina HiSeq 2000 using a single-read, 100 base read length (SR100) sequencing strategy.

### RNA-seq data processing and analysis

Image analysis and base calling were performed with Illumina’s Real Time Analysis v1.12 software. Files were demultiplexed of indexed reads and generated in FASTQ format using Illumina’s CASAVA v1.8. software. Reads were removed that did not pass Illumina’s base call quality threshold. Reads were aligned to mouse genome build mm9, using TopHat 1.4 (Trapnell et al., 2009). SAMtools v0.1.18 (Li et al., 2009) was used to sort and index the TopHat alignments. The gene expression profiles of embryos were compared using the Bioconductor package edgeR. Sequencing data will be available at NCBI GEO.

Functional annotation enrichment analysis was performed by submitting Ensemble Gene IDs from Table S1 and Table S2 to DAVID analysis (Huang da et al., 2009).

## Results

### Zebrafish mzpbx2;pbx4 mutant embryos show impaired activation of fast muscle differentiation

In zebrafish, the main *pbx* genes expressed during early embryonic myogenesis stages are *pbx2* and *pbx4* (Farr et al., 2018; Waskiewicz et al., 2002). Our previous studies of Pbx functions in zebrafish skeletal muscle development used antisense morpholinos to knock down both *pbx2* and *pbx4* expression, and we showed that these *pbx2-MO;pbx4-MO* embryos have delayed activation and reduced expression of fast muscle genes (Maves et al., 2007; Yao et al., 2013). Here, we have used CRISPR to generate a loss of function allele of zebrafish *pbx2* (Materials and Methods; Supplemental Figure 1). We find that *pbx2^-/-^* fish are homozygous viable as adults, similar to *Pbx2^-/-^* mice (Selleri et al., 2004). We crossed *pbx2* mutant fish with the established *pbx4* mutant strain (Pöpperl et al., 2000) to generate a *pbx2^-/-^;pbx4^+/-^* viable strain. Incrossing these fish generated *mzpbx2^-/-^;pbx4^-/-^* mutant embryos, thereby removing the maternal contribution of *pbx2* expression (Waskiewicz et al., 2002) in addition to the zygotic functions of *pbx2* and *pbx4*. We find that *mzpbx2^-/-^;pbx4^-/-^* embryos almost completely lack *krox20* expression in the hindbrain (Fig. 1), as expected for zebrafish embryos lacking Pbx proteins (Waskiewicz et al., 2002). We examined fast and slow muscle development in *mzpbx2^-/-^;pbx4^-/-^* embryos using RNA in situ hybridization (Fig. 1). As controls, we simultaneously collected embryos from incrosses of *pbx4^+/-^* fish. We observe that the fast muscle genes *atp2a1*, *myl1*, and *mylpfa*, as well as *myog*, show reduced expression in *mzpbx2^-/-^;pbx4^-/-^* embryos compared to somite-stage matched controls (Fig. 1). Trunk skeletal muscle expression of the slow muscle gene *smyhc1* appears largely unaffected in *mzpbx2^-/-^;pbx4^-/-^* embryos (Fig. 1). These results corroborate our previously described findings in *pbx2-MO;pbx4-MO* embryos (Maves et al., 2007; Yao et al., 2013). We observed no detectable effect on muscle development in *mzpbx2^-/-^* embryos and only subtle, variable effects on fast muscle gene expression in *pbx4^-/-^* embryos (data not shown). These results confirm that Pbx proteins are needed for the proper activation of fast muscle differentiation genes in zebrafish embryogenesis.

**Fig. 1.**
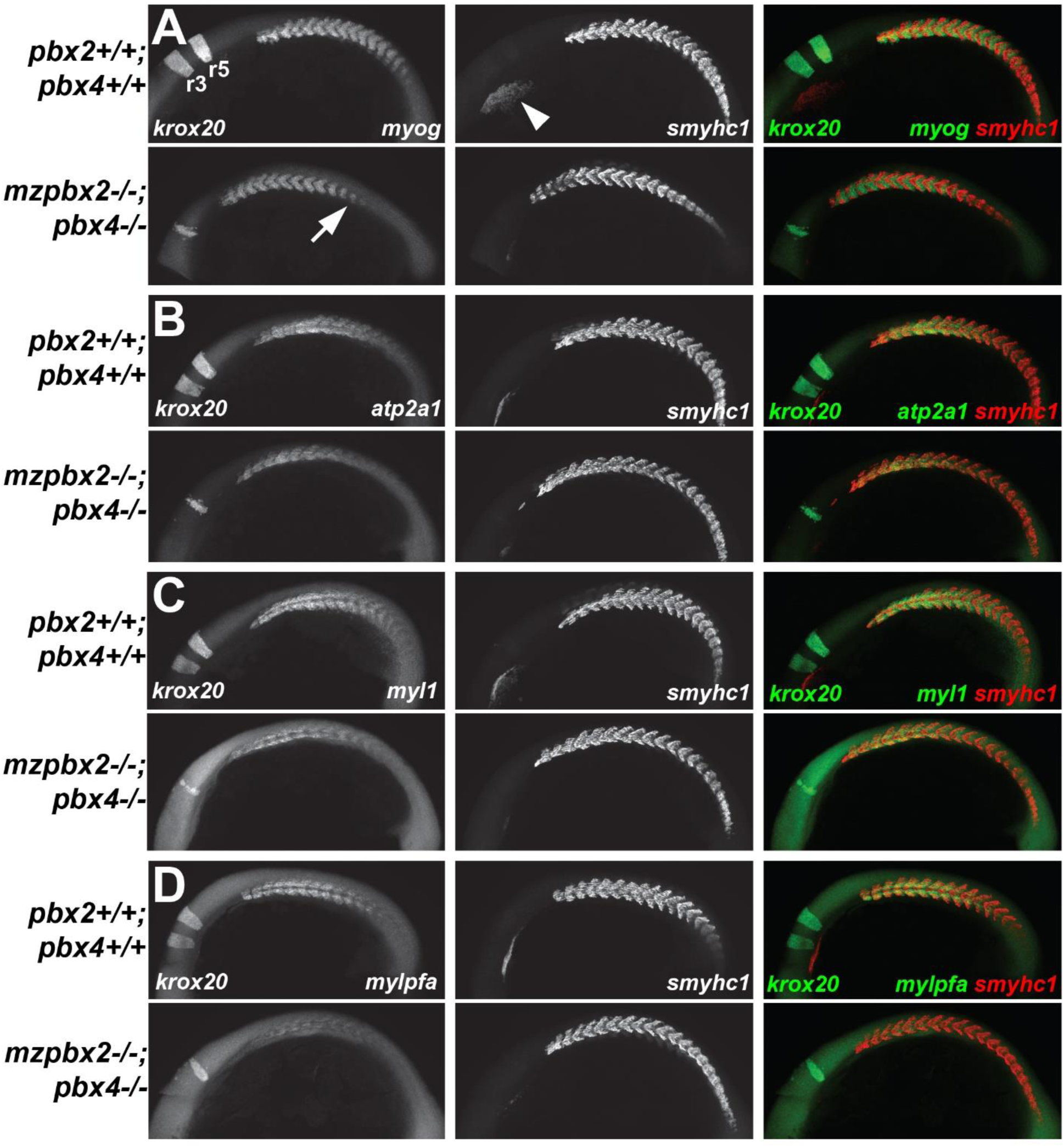
Fast muscle differentiation genes are downregulated in *mzpbx2^-/-^;pbx4^-/-^* zebrafish embryos relative to controls at 18 somite stage/18 hours post fertilization (hpf). (A-D) Whole mount RNA in situ hybridizations in 18 hpf embryos. Top panels of each set show control *pbx2^+/+^;pbx4^+/+^* embryos. Bottom panels of each set show somite-stage matched *mzpbx2^-/-^;pbx4^-/-^* embryos. Lateral views show anterior to the left and dorsal at the top. N=10-16 control and 3-5 mutant embryos for each condition. (A) Arrow in A points to reduced *myog* expression in *mzpbx2^-/-^;pbx4^-/-^* embryo. (B-D) *atp2a1*, *myl1*, and *mylpfa* appear generally reduced in *mzpbx2^-/-^;pbx4^-/-^* embryos. *krox20* is expressed in hindbrain rhombomeres 3 and 5 (r3, r5 in A) and is downregulated in *mzpbx2^-/-^;pbx4^-/-^* embryos. *smyhc1* expression in the developing heart (arrowhead in A) is downregulated in *mzpbx2^-/-^;pbx4^-/-^* embryos.

### Loss of Pbx1 has little effect on early mouse muscle differentiation

In mouse, previous studies have shown that *Pbx1* is expressed broadly in embryonic somites and is required for multiple developmental programs (Capellini et al., 2006; Selleri et al., 2001; Selleri et al., 2019). To investigate the requirements for Pbx factors in mouse muscle differentiation, we examined mouse embryos lacking Pbx1 (*Pbx1^-/-^*; Selleri et al., 2001). Using qRT-PCR, we find no significant effects on muscle fiber-type markers in E9.5 *Pbx1^-/-^* embryos (Fig. 2A). In E11.5 *Pbx1^-/-^* embryos, we find downregulation of *Myog* expression and upregulation of the slow muscle gene *Myh7*, but no significant changes in the other muscle differentiation genes (Fig. 2B). Using RNA in situ hybridization, we observe some downregulation of *Myog* expression in *Pbx1^-/-^* embryos at E11.5, consistent with our qRT-PCR results (Fig. 2C-D). These results suggest that loss of Pbx1 alone has subtle effects on early mouse muscle differentiation.

**Fig. 2.**
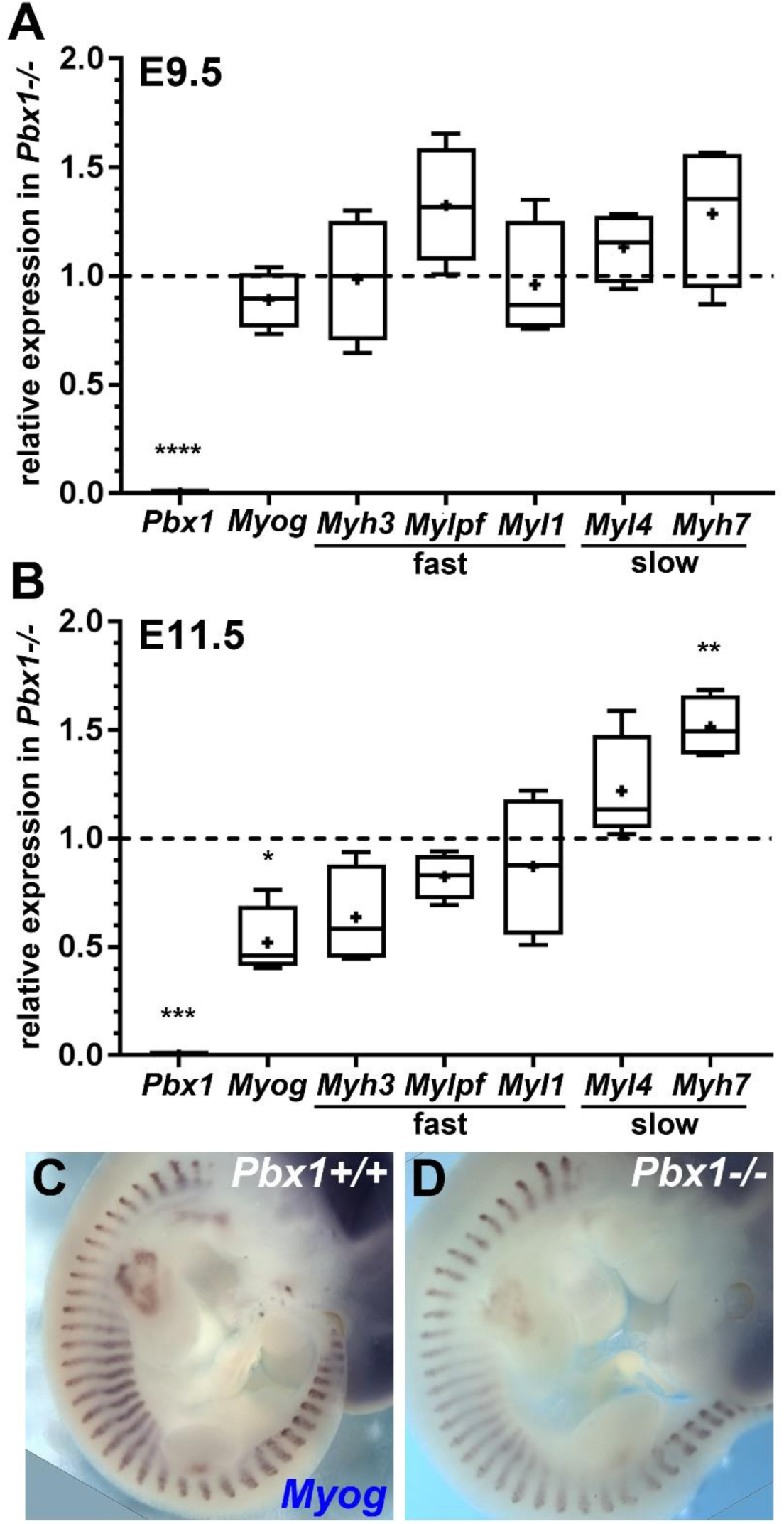
Loss of Pbx1 has little effect on early mouse muscle differentiation. (A-B) qRT-PCR analysis of muscle gene expression in (A) E9.5 and (B) E11.5 *Pbx1^-/-^* embryos relative to control *Pbx1^+/+^* embryos. *Myh3*, *Mylpf*, and *Myl1* are fast muscle genes. *Myl4* and *Myh7* are slow muscle genes. *Pbx1* expression is included as a control gene. Expression levels are normalized to the gene *Odc1*. For each gene, the expression level in *Pbx1^+/+^* embryos is set to 1 (dashed line in graphs). Box plots show the distribution of the normalized expression level (2^-ΔΔCt^) of four control/mutant sibling pairs for each gene. The boxes extend from the 25th to 75th percentiles, the whiskers are at the minimum and maximum, the bar within the box represents the median, and the cross represents the mean. Statistics are described in Materials and Methods. * P<0.02, ** P<0.004, *** P<0.0008, **** P<0.0001. (C-D) Whole mount *Myog* RNA in situ hybridization in (C) *Pbx1^+/+^* and (D) *Pbx1^-/-^* E11.5 embryos. N=3.

### Mouse Pbx1 and Pbx2 are both expressed in embryonic skeletal muscle precursors

Previous studies have shown that both *Pbx1* and *Pbx2* mRNAs are expressed broadly in mouse somites at E8.5 (Capellini et al., 2006). We thus wanted to determine whether Pbx1 and Pbx2 proteins are expressed in mouse embryo somites, which contain the primary myotome, the precursors of early mouse muscle (Biressi et al., 2007). Using antibody staining, we observe broad somitic expression of Pbx1 and Pbx2 at E11.5, (Fig. 3A,C), and they both show nuclear localization (Fig. 3B,D). Pbx1 and Pbx2 show overlap with Myf5 and Myod1 (Fig. 3A-D), which both mark myogenic cells in the somites (Comai et al., 2014; Relaix et al., 2005). Thus, both Pbx1 and Pbx2 are expressed in embryonic mouse myogenic progenitors.

**Fig. 3.**
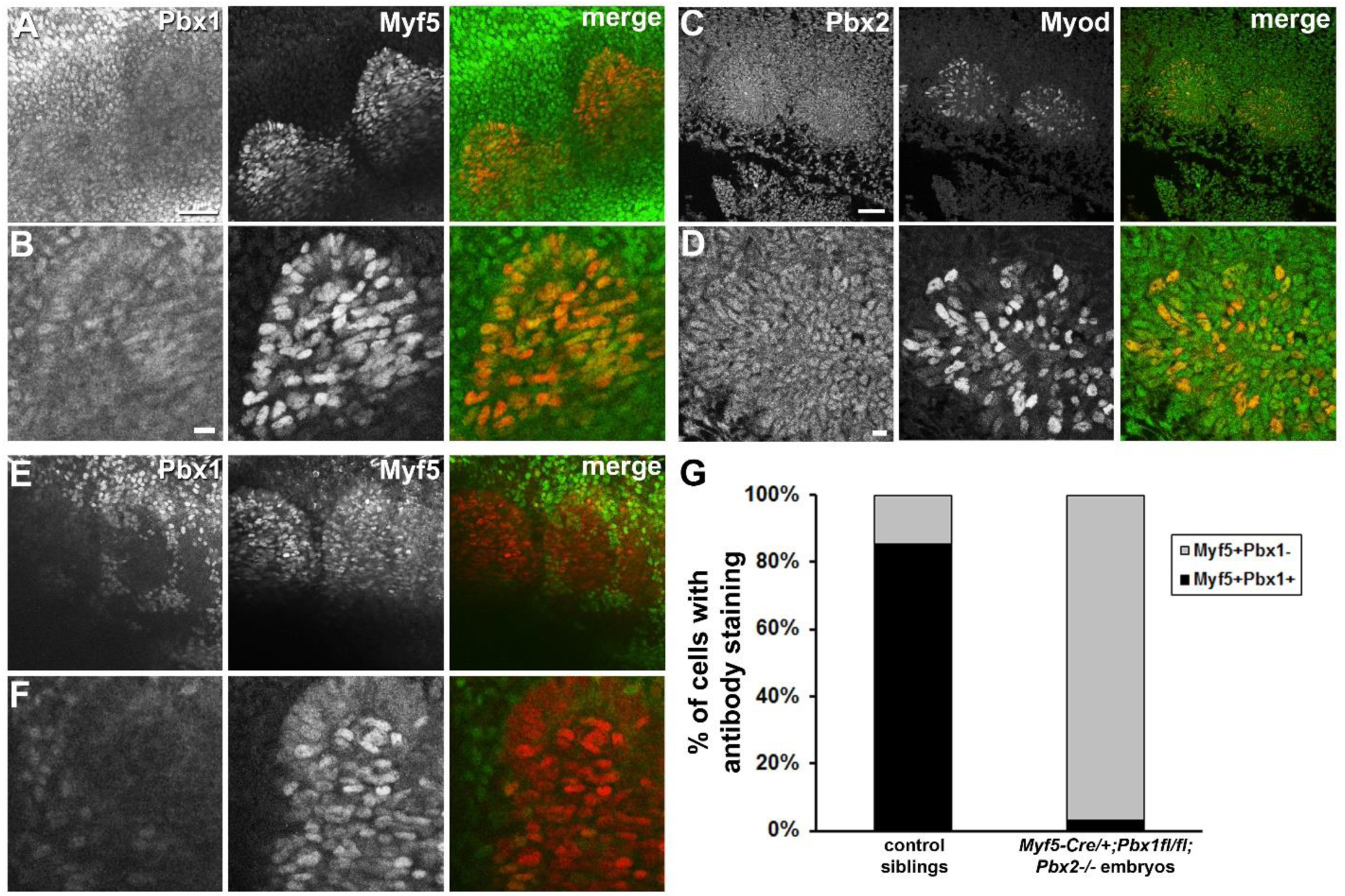
Pbx1 and Pbx2 expression in mouse embryo somites. (A-B) E11.5 mouse embryo whole mount stained with anti-Pbx1b and anti-Myf5. A, low magnification, scale bar=20 µm. B, high magnification of A, scale bar=20 µm. (C-D) E11.5 mouse embryo sagittal section stained for anti-Pbx2 and anti-Myod1. C, low magnification, scale bar=10 µm. D, high magnification of C, scale bar=10 µm. (E-F) E11.5 *Myf5^Cre/+^;Pbx1^fl/fl^* embryo stained with anti-Pbx1b and anti-Myf5. E, low magnification, scale bar as in A. F, high magnification of E, scale bar as in B. Rostral is to the left. (G) Quantification of Myf5 and Pbx1b co-expressing cells (Myf5+Pbx1+) in E11.5 control embryos and *Myf5^Cre/+^;Pbx1^fl/fl^;Pbx2^-/-^* mutant embryos.

### Conditional loss of Pbx1 and Pbx2 reveals requirements for mouse Pbx proteins in early fast muscle differentiation

Because several studies have shown that Pbx1 and Pbx2 have compensatory and overlapping functions (Capellini et al., 2006; Ferretti et al., 2011; Koss et al., 2012), we next wanted to address requirements for both Pbx1 and Pbx2 in early mouse muscle differentiation. *Pbx2^-/-^* mice are viable (Selleri et al., 2004). *Pbx1^-/-^;Pbx2^-/-^* compound constitutive mutant embryos have multiple organogenesis defects and die by E10.5 (Capellini et al., 2006), precluding an examination of early muscle differentiation. Therefore, we decided to examine conditional loss of *Pbx1* on a *Pbx2^-/-^* background (Ferretti et al., 2011; Koss et al., 2012). We used the *Myf5-Cre* deleter strain (Gensch et al., 2008; Tallquist et al., 2000) in combination with the conditional *Pbx1* allele (*Pbx1^fl^*; Koss et al, 2012) to remove Pbx1 from early myogenic cells. In *Myf5^Cre/+^;Pbx1^fl/fl^;Pbx2^-/-^* embryos, we used antibody staining to show loss of Pbx1 in Myf5-expressing cells in the somites (Fig. 3E-G), confirming efficient Cre-mediated inactivation of Pbx1. We find that *Myf5^Cre/+^;Pbx1^fl/fl^* mice are viable and exhibit kyphosis as adults (data not shown), indicating a possible muscle weakness phenotype. We crossed *Myf5^Cre/+^;Pbx1^fl/fl^* mice with the *Pbx2^-/-^* line to remove both Pbx1 and Pbx2 from muscle precursors. In crosses of *Pbx1^fl/fl^;Pbx2^-/-^* mice with *Myf5^Cre/+^;Pbx1^fl/+^;Pbx2^-/-^* mice, we observe expected Mendelian ratios for conditional double mutant *Myf5^Cre/+^;Pbx1^fl/fl^;Pbx2^-/-^* embryos at E11.5 and E17.5, but we obtain negligible numbers of *Myf5^Cre/+^;Pbx1^fl/fl^;Pbx2^-/-^* pups at postnatal day 7 (Table 2). From multiple litters, we were able to identify 3 dead P0-P1 pups in the cage bedding, that, upon genotyping were *Myf5^Cre/+^;Pbx1^fl/fl^;Pbx2^-/-^*. These results indicate that *Myf5^Cre/+^;Pbx1^fl/fl^;Pbx2^-/-^* mutant animals likely die at or just after birth.

**Table 2.** Offspring of Pbx1^fl/fl^;Pbx2^-/-^ X Myf5^Cre/+^;Pbx1^fl/+^;Pbx2^-/-^Mice

| Genotype | Frequencies at E11.5 |  |  | Frequencies at E17.5 |  |  | Frequencies at P7 |  |  |
| --- | --- | --- | --- | --- | --- | --- | --- | --- | --- |
|  | Observed | Expected | P | Observed | Expected | P | Observed | Expected | P |
| *Myf5<sup>Cre/+</sup>;*<br>*Pbx1<sup>fl/fl</sup>;Pbx2<sup>-/-</sup>* | 25 | 22.25 | 0.902 | 7 | 12 | 0.227 | 2 | 20.25 | 4.335<br>E-05 |
| *Myf5<sup>Cre/+</sup>;*<br>*Pbx1<sup>fl/+</sup>;Pbx2<sup>-/-</sup>* | 20 | 22.25 |  | 13 | 12 |  | 29 | 20.25 |  |
| *Pbx1<sup>fl/fl</sup>;Pbx2<sup>-/-</sup>* | 22 | 22.25 |  | 17 | 12 |  | 27 | 20.25 |  |
| *Pbx1<sup>fl/+</sup>;Pbx2<sup>-/-</sup>* | 22 | 22.25 |  | 11 | 12 |  | 23 | 20.25 |  |

P values calculated using a Chi-square test.

To determine the effects of conditional loss of Pbx1 and Pbx2 on early muscle differentiation, we collected RNA from whole *Myf5^Cre/+^;Pbx1^fl/fl^;Pbx2^-/-^* embryos and their sibling controls at E11.5 and performed RNA-seq. Using the cut-offs of 1.2 fold change (FC) and P<0.05, we identified 229 genes down-regulated and 144 genes up-regulated in *Myf5^Cre/+^;Pbx1^fl/fl^;Pbx2^-/-^* mutants (Supplementary Tables S1-S2). DAVID bioinformatics analysis (Huang da et al., 2009) revealed significant enrichment of multiple functional annotation terms related to muscle development and differentiation in the genes down-regulated in *Myf5^Cre/+^;Pbx1^fl/fl^;Pbx2^-/-^* mutant embryos (Fig. 4A). For the up-regulated gene set, DAVID analysis did not identify any functional annotation terms with significant enrichment following multiple testing correction (not shown). Within the gene set showing downregulation in *Myf5^Cre/+^;Pbx1^fl/fl^;Pbx2^-/-^* embryos (Table S1), we noticed the presence of genes associated with fast-twitch muscle differentiation, including genes that we showed to be Pbx-dependent in zebrafish, such as *Mylpf* (Fig. 1; Maves et al., 2007; Yao et al., 2013). We directly examined muscle fiber-type-associated genes (Schiaffino and Reggiani, 2011) in the RNA-seq data and found 11/13 fast muscle genes with significant down-regulation (FC > 1.2, P > 0.05) in *Myf5^Cre/+^;Pbx1^fl/fl^;Pbx2^-/-^* embryos, whereas 9/14 slow muscle genes showed either up-regulation or no significant change (Fig. 4B). We used qRT-PCR to validate some of these genes on an additional set of *Myf5^Cre/+^;Pbx1^fl/fl^;Pbx2^-/-^* embryos and their control siblings and found that, indeed, fast muscle genes showed significant down-regulation, whereas slow muscle genes generally are unaffected (Fig. 4C).

**Fig. 4.**
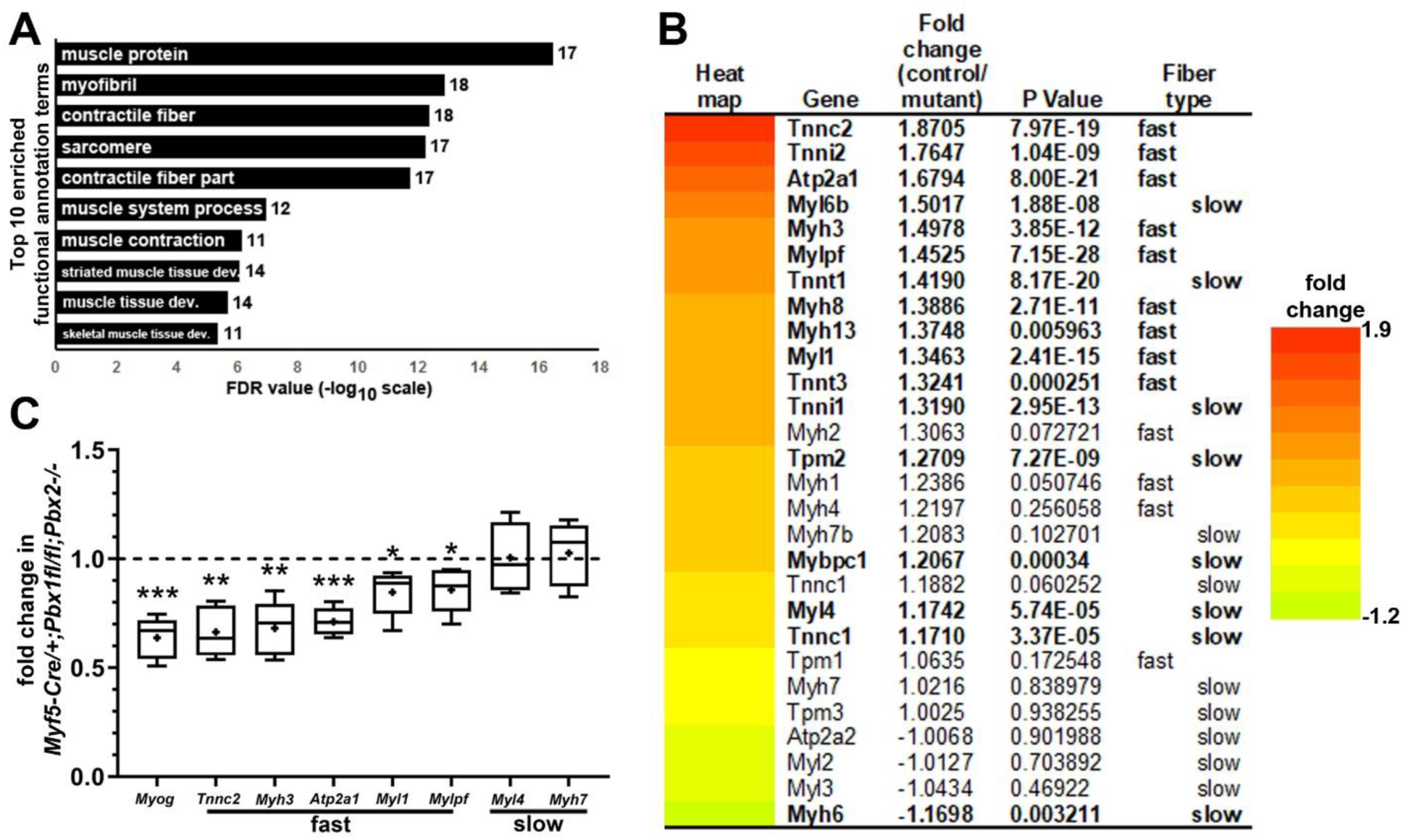
*Pbx1* and *Pbx2* are needed for mouse embryonic fast muscle gene expression. (A) Functional annotation terms associated with genes downregulated (Table S1) in E11.5 *Myf5^Cre/+^;Pbx1^fl/fl^;Pbx2^-/-^* embryos relative to *Pbx1^fl/+^;Pbx2^-/-^* control siblings. Numbers next to each bar indicate number of genes associated with each term. (B) Muscle fiber type-associated gene expression from RNA-seq in mutant *Myf5^Cre/+^;Pbx1^fl/fl^;Pbx2^-/-^* E11.5 embryos relative to *Pbx1^fl/+^;Pbx2^-/-^* control siblings. Genes showing significant changes are in bold. Color bar legend illustrates range of fold changes. (C) qRT-PCR analysis of muscle gene expression in E11.5 *Myf5^Cre/+^;Pbx1^fl/fl^;Pbx2^-/-^* embryos relative to control siblings. Control siblings consisted of *Pbx1^fl/+^;Pbx2^-/-^* (N=1) or *Myf5^Cre/+^;Pbx1^fl/+^;Pbx2^-/-^* (N=4) embryos. *Tnnc2*, *Myh3*, *Atp2a1*, *Myl1* and *Mylpf* are fast muscle genes. *Myl4* and *Myh7* are slow muscle genes. Expression levels are normalized to the gene *Odc1*. For each gene, the expression level in control sibling embryos is set to 1 (dashed line). Box plots show the distribution of the normalized expression level (2^-ΔΔCt^) of five control/mutant sibling pairs for each gene. The boxes extend from the 25th to 75th percentiles, the whiskers are at the minimum and maximum, the bar within the box represents the median, and the cross represents the mean. Statistics are described in Materials and Methods. * P<0.05, ** P<0.01, *** P<0.003.

We next validated the results from our RNA-seq and qRT-PCR analyses by performing RNA in situ hybridization experiments. We tested *Myog* and some of the top down-regulated fast muscle genes (*Tnnc2*, *Myh3*, *Atp2a1*, *Myl1*, and *Mylpf*) by RNA in situ at E11.5 (Fig. 5). We observe that these fast muscle genes show reduced expression in *Myf5^Cre/+^;Pbx1^fl/fl^;Pbx2^-/-^* embryos compared to somite-stage matched control siblings (Fig. 5). Slow muscle genes *Myl4*, *Myh7*, and *Myh6* appear largely unaffected in *Myf5^Cre/+^;Pbx1^fl/fl^;Pbx2^-/-^* embryos at E11.5 (Fig. 5). In addition, *Myh3*, *Myl4*, *Myh7*, and *Myh6* have strong heart expression that appears unaffected in *Myf5^Cre/+^;Pbx1^fl/fl^;Pbx2^-/-^* embryos (Fig. 5). These findings are consistent with our RNA-seq and qRT-PCR analyses and establish that Pbx proteins are needed for the proper activation of a program of fast muscle differentiation genes in mouse embryogenesis. Taken together, these results confirm that Pbx proteins are needed for the proper activation of a program of fast muscle differentiation genes in both mouse and zebrafish embryogenesis.

**Fig. 5.**
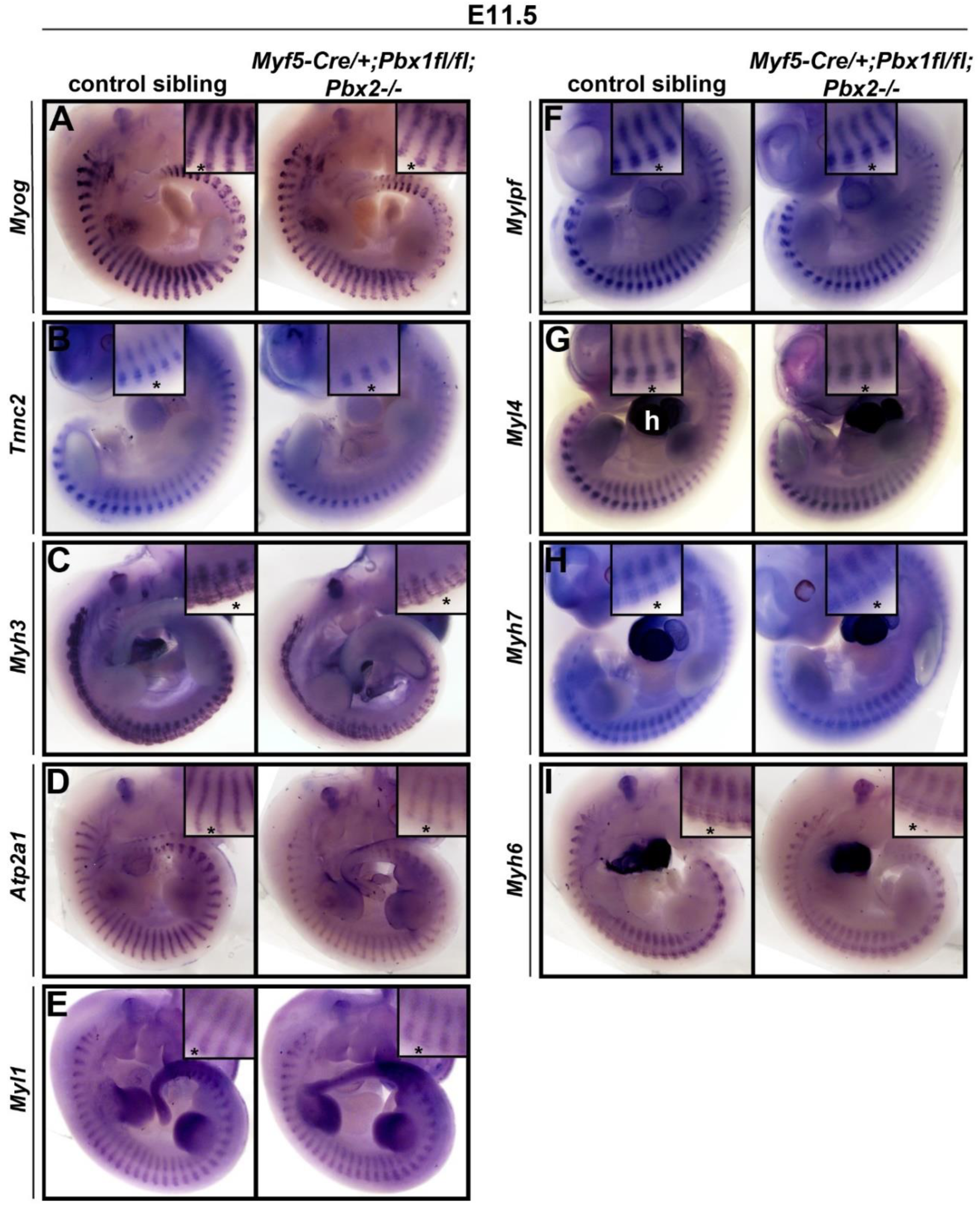
Fast muscle differentiation genes are downregulated in *Myf5^Cre/+^;Pbx1^fl/fl^;Pbx2^-/-^* embryos relative to control siblings at E11.5. (A-I) Whole mount RNA in situ hybridizations in E11.5 embryos. Left column of each panel shows control embryos, and right column of each panel shows somite-stage matched *Myf5^Cre/+^;Pbx1^fl/fl^;Pbx2^-/-^* embryos. The pictured control embryos for *Myog*, *Tnnc2*, *Myh3*, *Mylpf*, *Myl4*, *Myh7*, and *Myh6* are *Myf5^Cre/+^;Pbx1^fl/+^;Pbx2^-/-^*. The pictured control embryo for *Atp2a1* is *Pbx1^fl/fl^;Pbx2^-/-^*, and the control for *Myl1* is *Pbx1^fl/+^;Pbx2^-/-^*. No differences in gene expression were noted among controls of different genotypes. Insets show 2X zoom views at the level of somite 17 (* in each panel). N=3-5 control and 3-5 mutant embryos for each condition. *Myh3*, *Myl4*, *Myh7*, and *Myh6* have strong heart expression (h in C) that appears unaffected in *Myf5^Cre/+^;Pbx1^fl/fl^;Pbx2^-/-^* embryos.

## Discussion

Here we show evolutionary conservation of requirements for Pbx proteins in mammalian and zebrafish embryonic fast-twitch muscle differentiation. These results establish that Pbx homeodomain proteins have conserved roles in the control of the activation of fast muscle differentiation genes in different species.

Because of the redundancy of, and early requirements for, Pbx proteins in mouse embryogenesis (reviewed in Selleri et al. 2019), we needed to make use of conditional loss-of-function in *Myf5^Cre/+^;Pbx1^fl/fl^;Pbx2^-/-^* compound mutants. A similar approach, using different conditional *Cre* lines, has previously been taken to study Pbx functions in craniofacial, spleen, and spinal cord motor neuron development and in adult neurogenesis (Ferretti et al., 2011; Grebbin et al., 2016; Hanley et al., 2016; Koss et al., 2012; Losa et al., 2018; Welsh et al., 2018). Based on their expression patterns, *Pbx1* and *Pbx2* are the *Pbx* genes most likely to be involved in mouse embryonic skeletal muscle development (Capellini et al., 2006; Di Giacomo et al., 2006; Wagner et al., 2001). Future studies could additionally include loss of *Pbx3* (Rhee et al., 2004) to determine whether this Pbx family member also functions in muscle development and fiber-type differentiation.

We chose the *Myf5-Cre* line to conditionally knock out *Pbx1* in early skeletal muscle development because *Myf5* is expressed early in developing somites in the presumptive primary myotome, prior to *Myod1* expression (Kassar-Duchossoy et al., 2004; Ott et al., 1991). The *Myf5-Cre* line we employed has been previously used to examine satellite cells in limb and extraocular muscles, to ablate embryonic myogenic cells, and to conditionally ablate genes in skeletal muscle (Comai et al., 2014; Gensch et al., 2008; Greschik et al., 2017; Kuang et al., 2007; Stuelsatz et al., 2014). We note that the *Myf5-Cre* allele may not be effecting a full deletion of *Pbx1* in all myoblasts (Fig. 3; Comai et al., 2014), possibly explaining our finding that slow muscle gene *Myh7* is upregulated in *Pbx1^-/-^* mouse embryos (Fig. 2B) but does not appear affected in *Myf5^Cre/+^;Pbx1^fl/fl^;Pbx2^-/-^* embryos (Fig. 4C). However, we observe similar defects in the activation of fast muscle differentiation following both conditional inactivation of *Pbx1* and *Pbx2* in mouse embryos and whole-animal *mzpbx2;pbx4* loss-of-function in zebrafish embryos. These results indicate that the *Myf5-Cre* line is enabling adequate *Pbx* gene inactivation, leading to a deeper understanding of the function of this family of homeodomain proteins in muscle development in different species.

The defects in activation of fast muscle differentiation genes that we observe in *Myf5^Cre/+^;Pbx1^fl/fl^;Pbx2^-/-^* mouse embryos are similar to those observed in *Six1;Six4* double mutant mouse embryos (Niro et al., 2010). In *Six1^-/-^;Six4^-/-^* mouse embryos, activation of fast muscle genes, including *Atp2a1*, *Myl1*, and *Tnnc2*, is inhibited, while slow muscle genes, such as *Myl4*, are still expressed (Niro et al., 2010). Six1 and Six4 proteins directly bind regulatory regions of fast muscle genes (Niro et al., 2010). In future studies, it will be of interest to examine potential genetic and biochemical interactions between Pbx proteins and Six proteins during muscle development. In addition, such further studies, using both zebrafish and mice, could help advance our understanding of the conserved and non-conserved roles for Pbx and Six factors in muscle development and fiber-type differentiation during vertebrate evolution.

In conclusion, we provide evidence for evolutionarily conserved regulation of embryonic fast muscle differentiation by Pbx factors. In future studies, we plan to determine the requirements for Pbx factors in later stages of muscle development and fiber-type differentiation in both zebrafish and mice. In addition, we plan to take advantage of our zebrafish and mouse mutant strains to examine, in developing embryos, the Pbx-dependent chromatin events in myogenesis and muscle differentiation. The goal will be to determine whether Pbx proteins are needed during skeletal muscle development to open chromatin, or to promote Myod1 binding or Myod1 activity. Ultimately, these studies will help us understand how broadly-expressed factors like Pbx help master regulatory factors like Myod1 activate their transcriptional targets in a temporally and spatially controlled manner to drive cell-type-specific differentiation programs in different species.

## Acknowledgements

We thank Peter Currie, Zhiyuan Gong, Phil Ingham, Heike Pöpperl, and Phil Soriano for generously providing reagents. The Zebrafish International Resource Center (supported by grants RR12546 and RR15402-01 from the NIH) provided cDNA clones. We thank the FHCRC Genomics Shared Resource for excellent technical assistance with the RNA-seq experiments. We thank the SCRI Aquatics Facility for zebrafish care. This work was supported by NIH 1R03AR057477 (L.M.), funded in part by the American Recovery and Reinvestment Act, NIH 1R03AR065760 (L.M.), the Seattle Children’s Myocardial Regeneration Initiative, by NIH 1R01AR45113 (S.J.T.), and by NIH RO1 DE024745 (L.S.).

---

**Supplemental Fig. 1.**
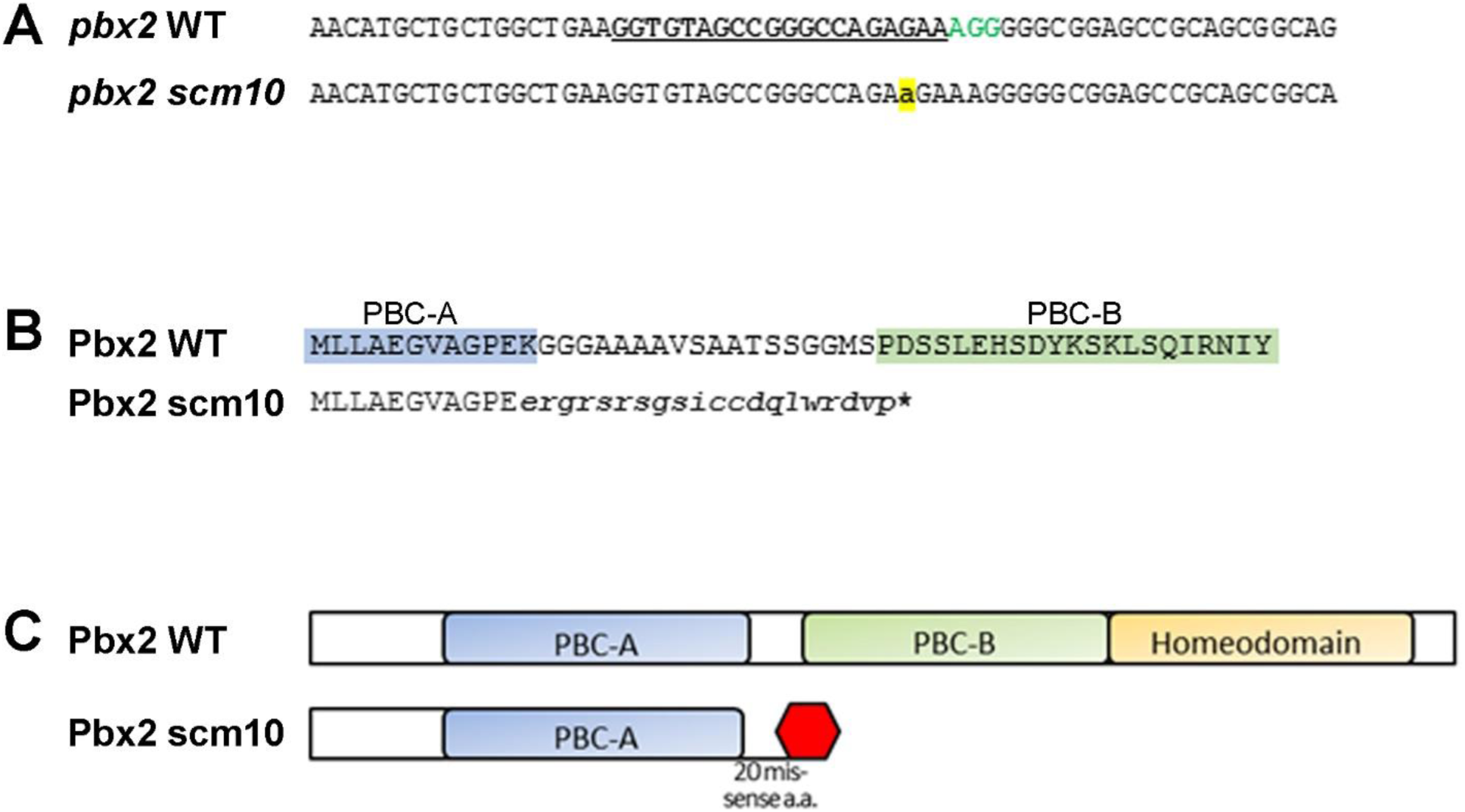
Generation of a zebrafish *pbx2* mutant allele with CRISPR/Cas9. (A) Top line: a portion of the wild-type (WT) genomic sequence of *pbx2* from the middle of Exon 3 showing the target sequence for the single-guide RNA (bold, underline) and the PAM sequence (green, bold). Bottom line: the sequence of the *pbx2^scm10^* allele with a single “A” nucleotide (highlighted) inserted 3bp upstream of the PAM. (B) Alignment of the conceptual translation of the *pbx2^scm10^* allele compared to wild-type Pbx2. Mis-sense amino acids are in lower case and the premature STOP codon is noted with an asterisk. The PBC-A and -B domains are conserved among the Pbx family. (C) Cartoon showing the domain structure of Pbx2. The scm10 mutation is predicted to produce a truncated product lacking the conserved PBC-B and Homeobox domains and thus is likely to be a loss of function allele.

**Table S1.**
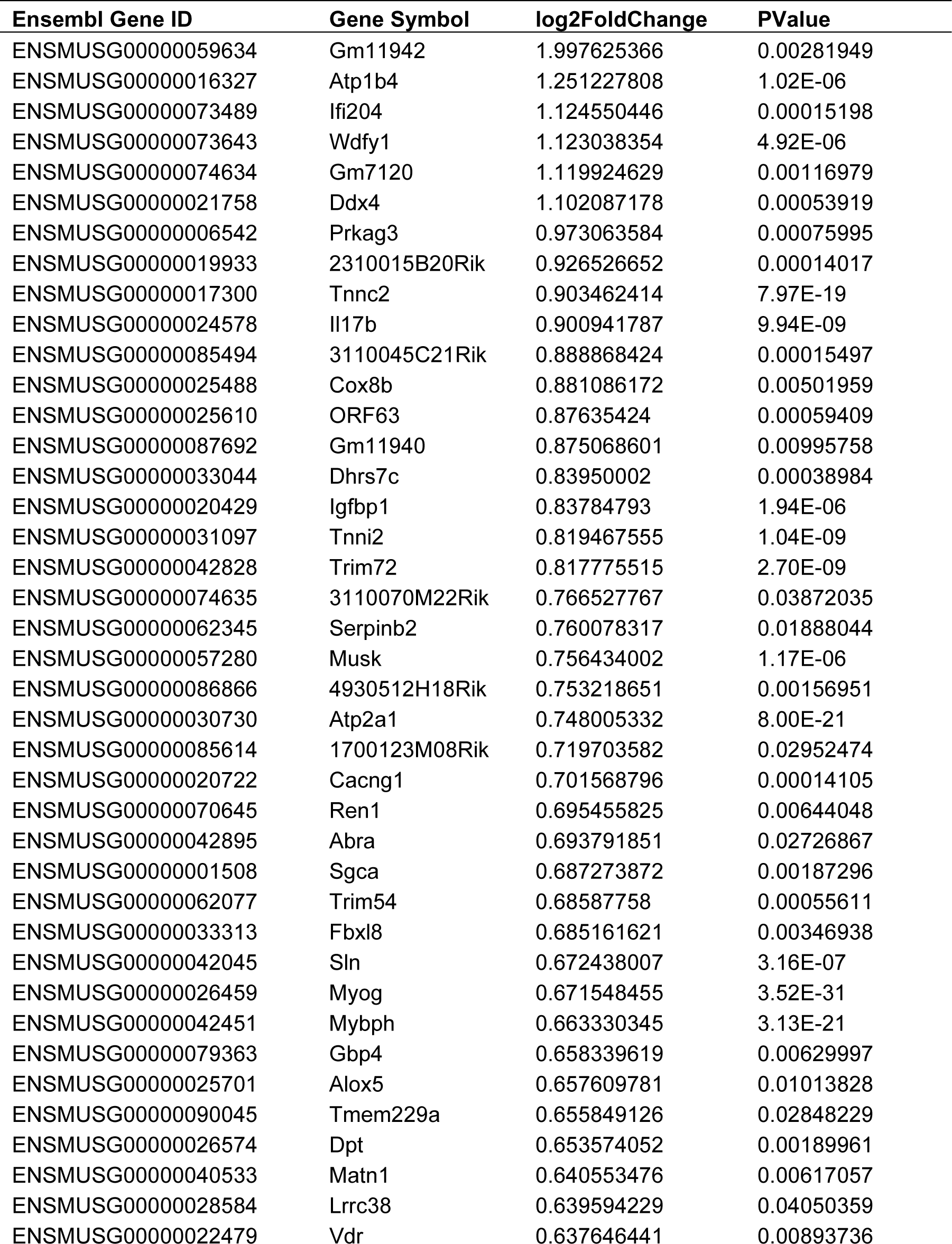
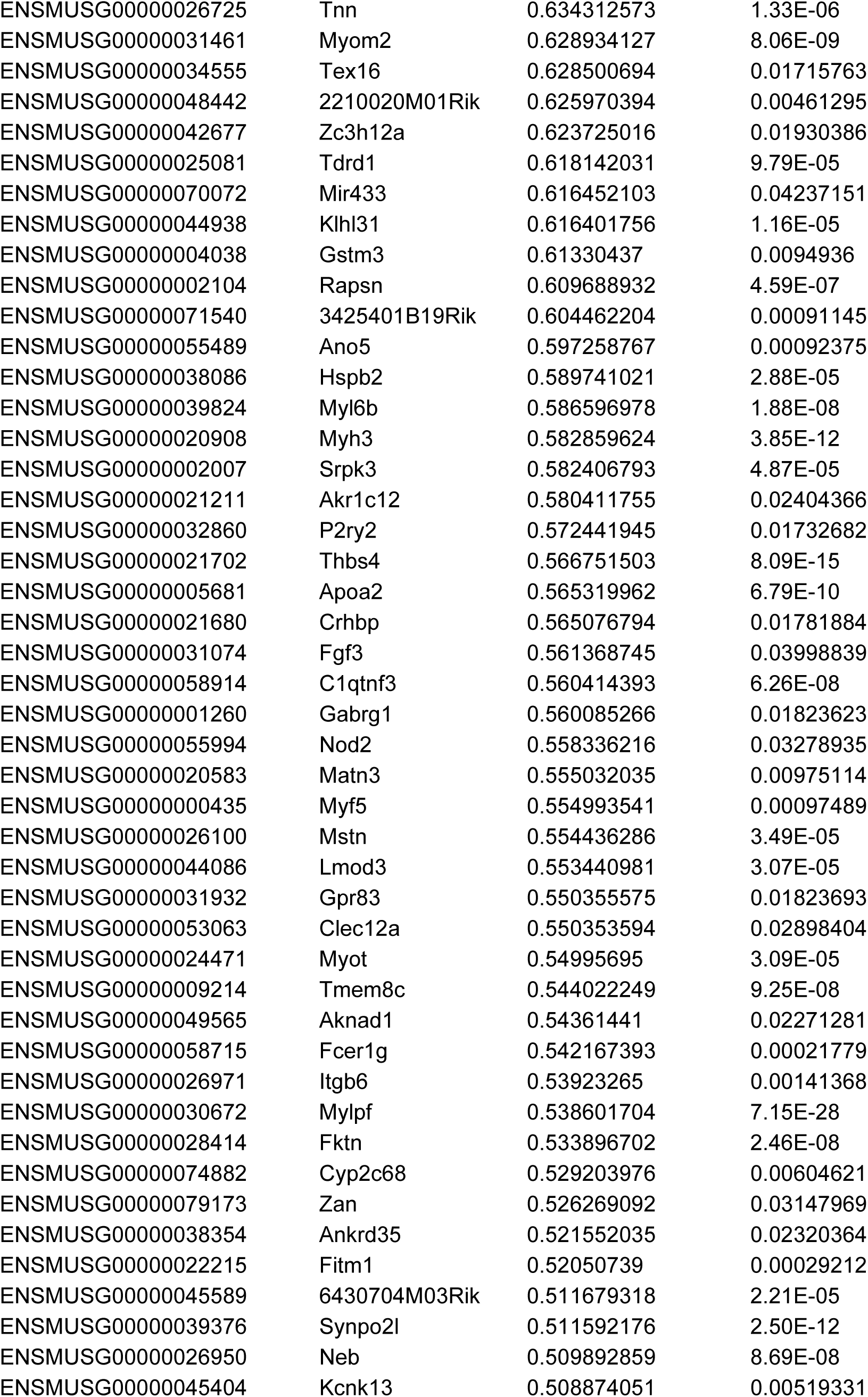
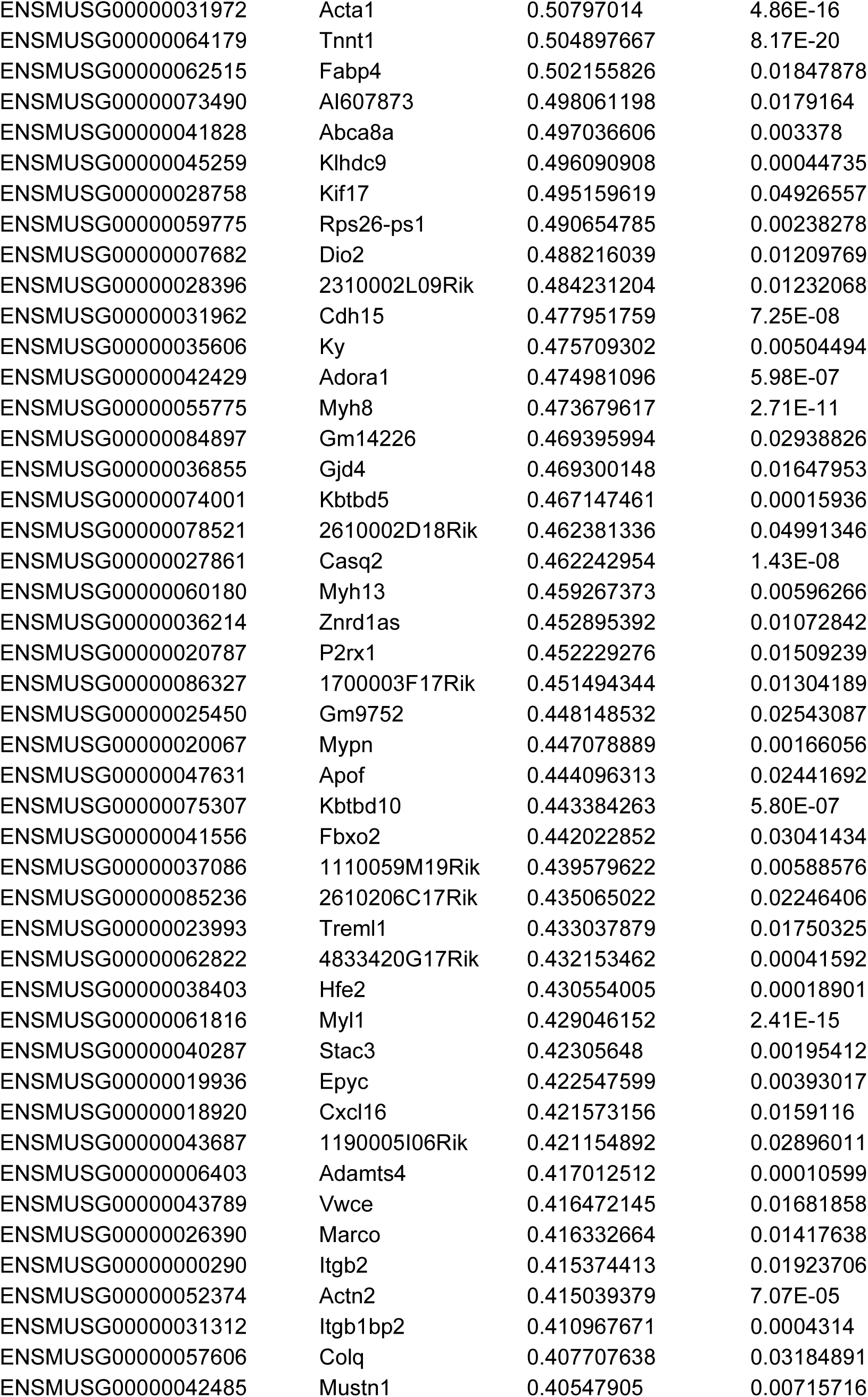
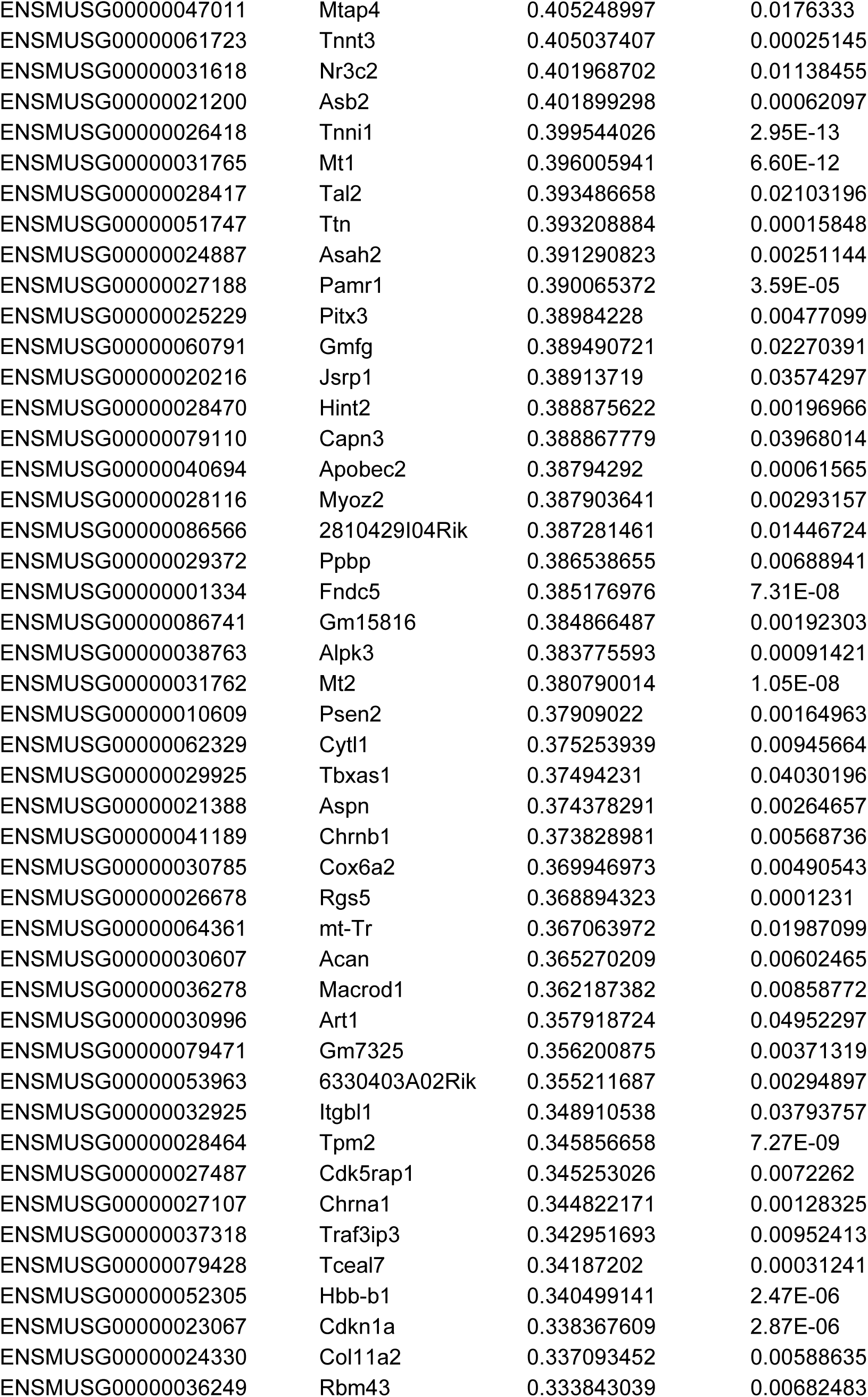
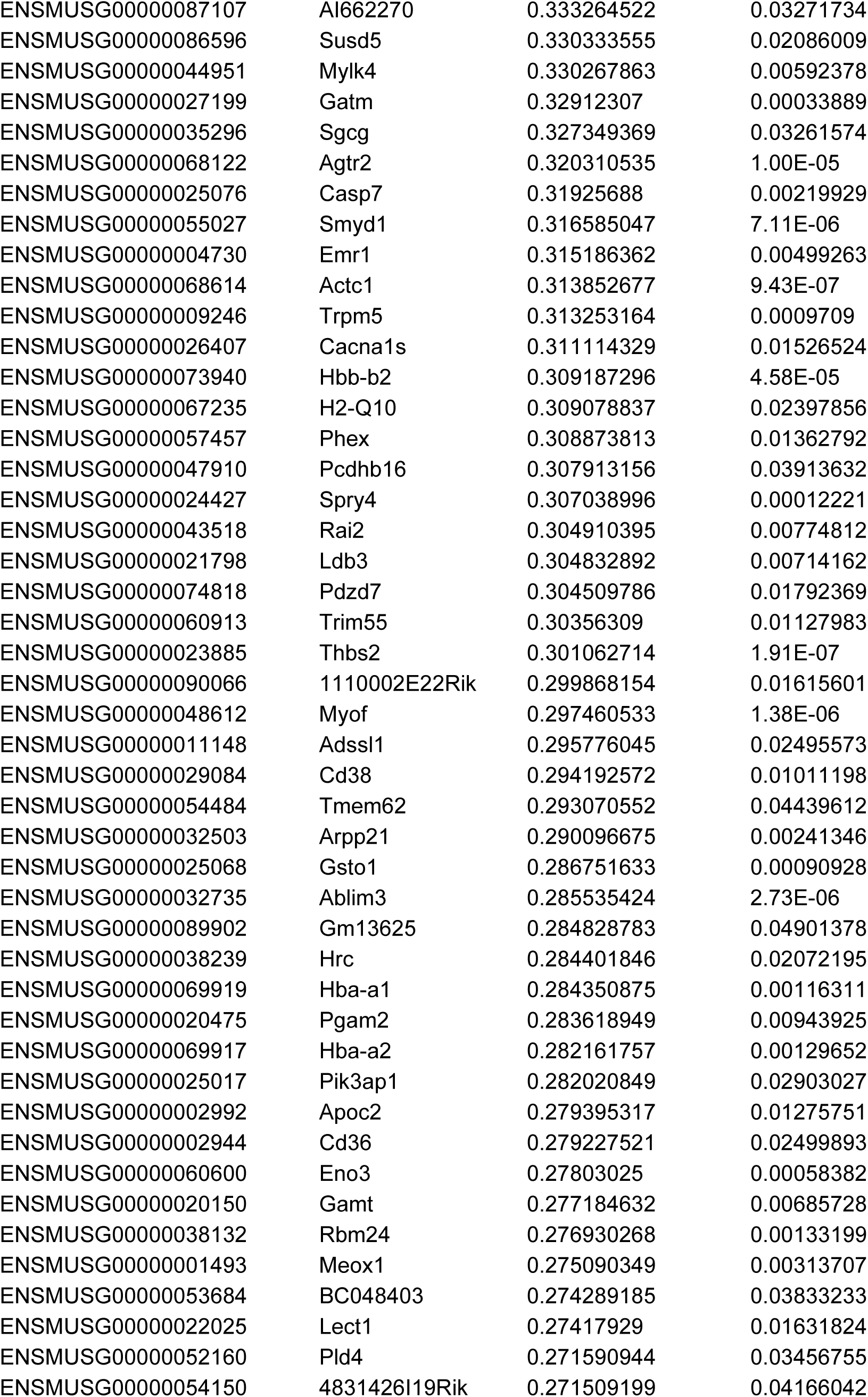
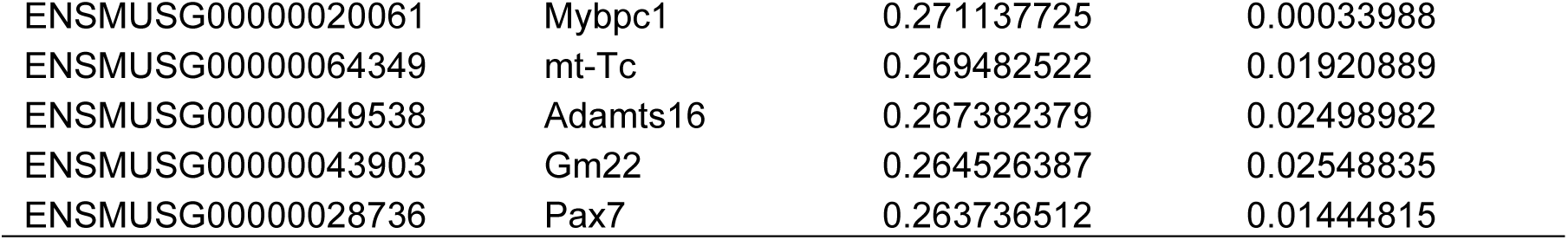
Genes downregulated in *Myf5^Cre/+^;Pbx1^fl/fl^;Pbx2^-/-^* E11.5 embryos (FC>1.2, P<0.05)

**Table S2.**
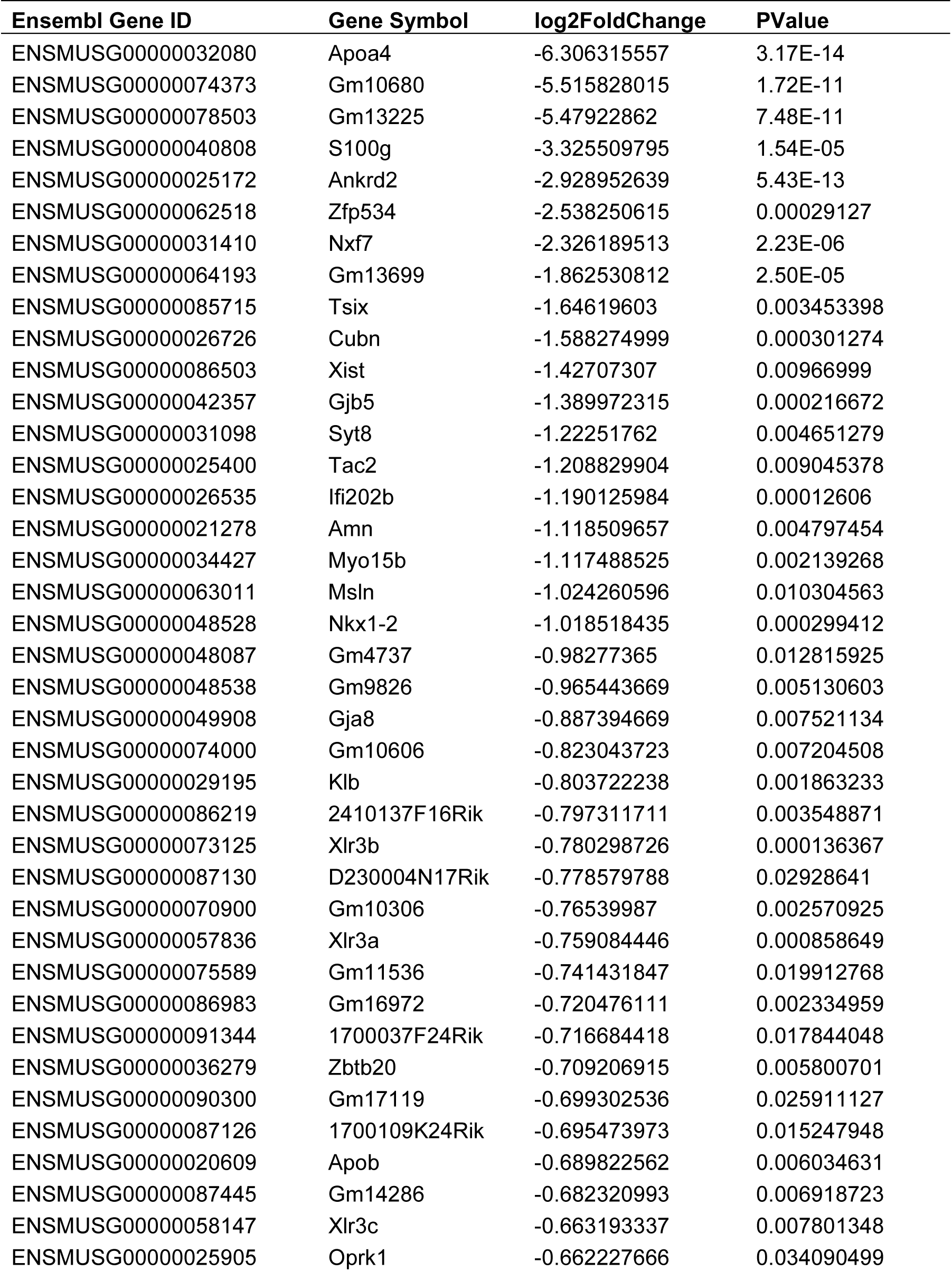
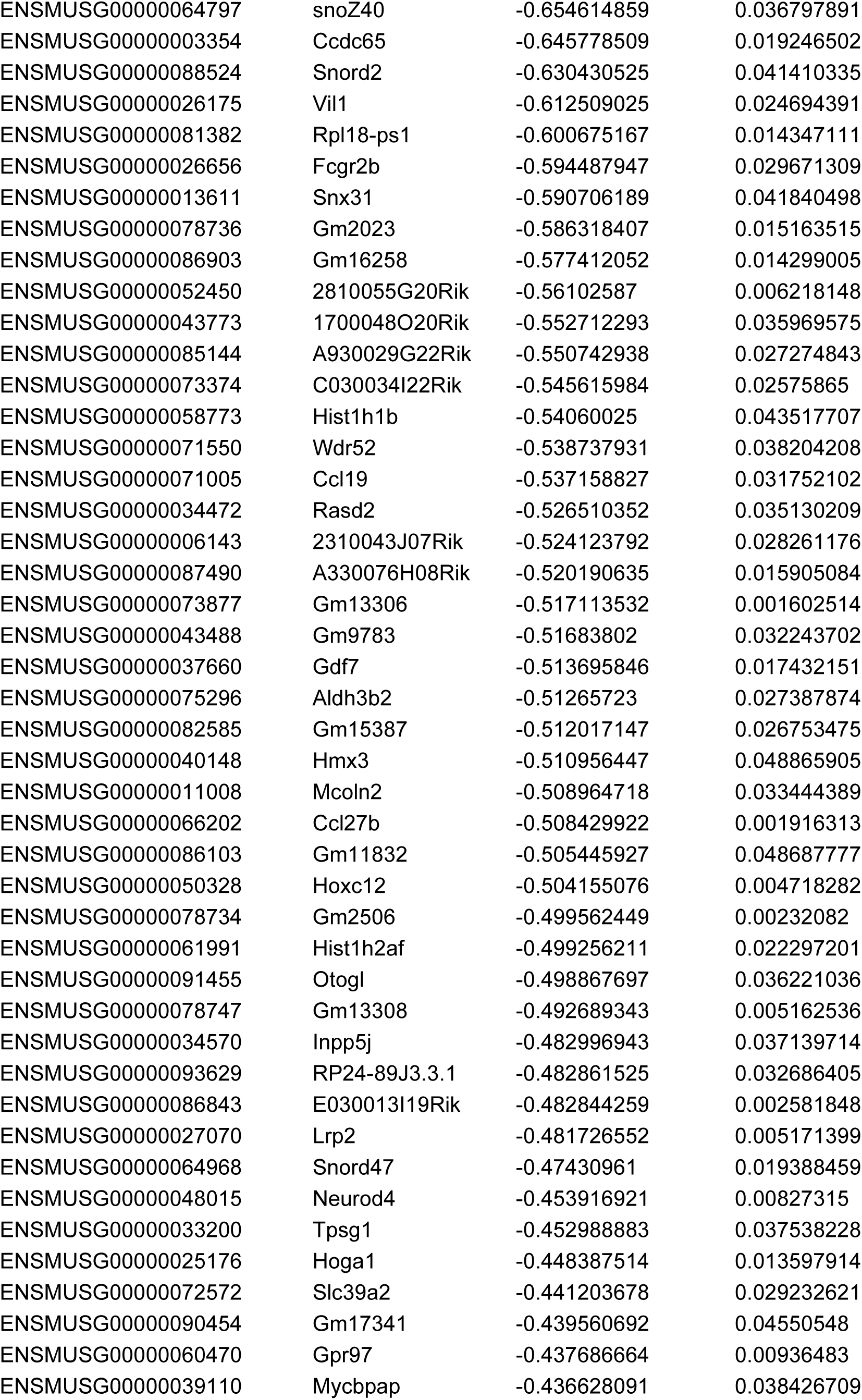
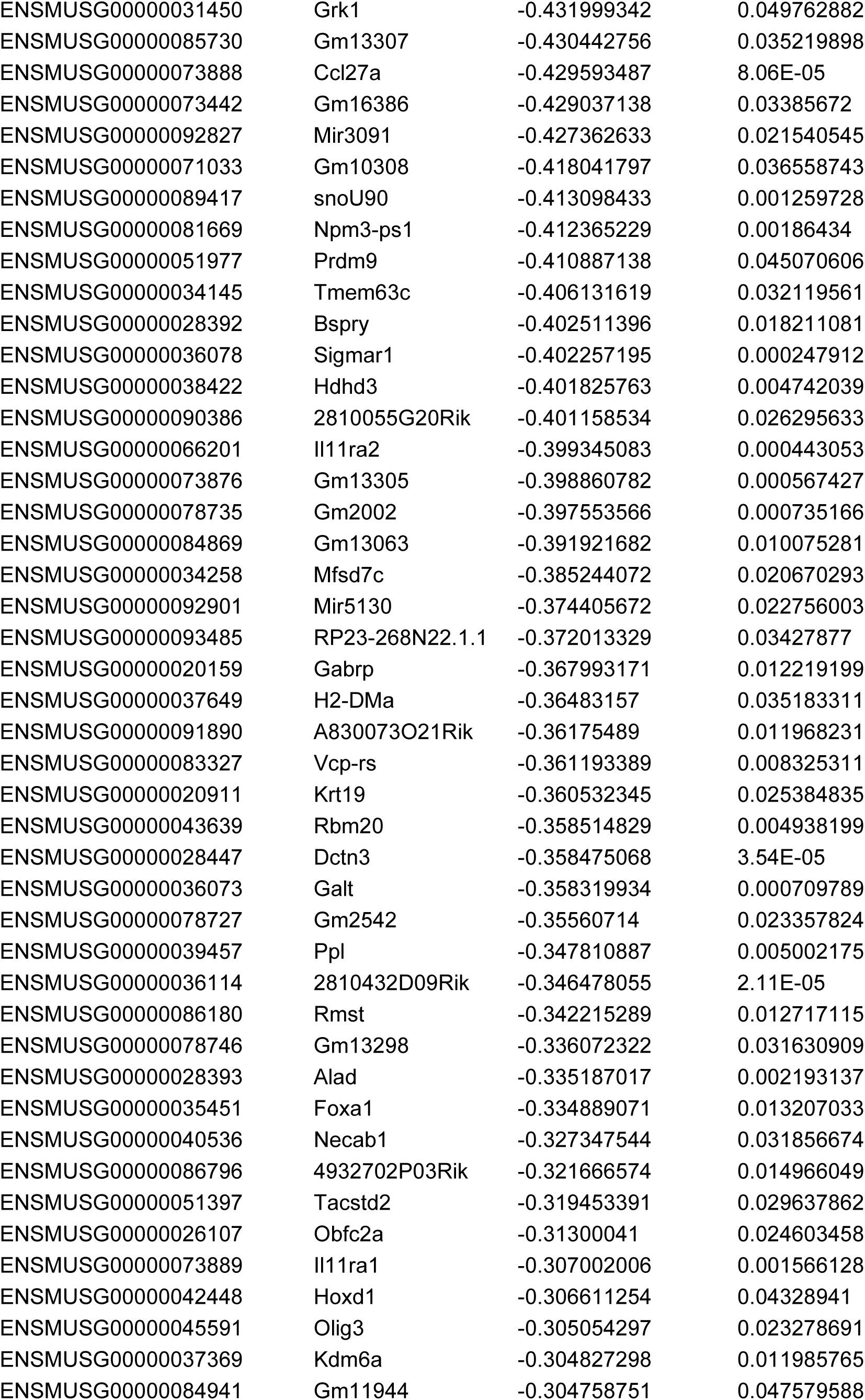
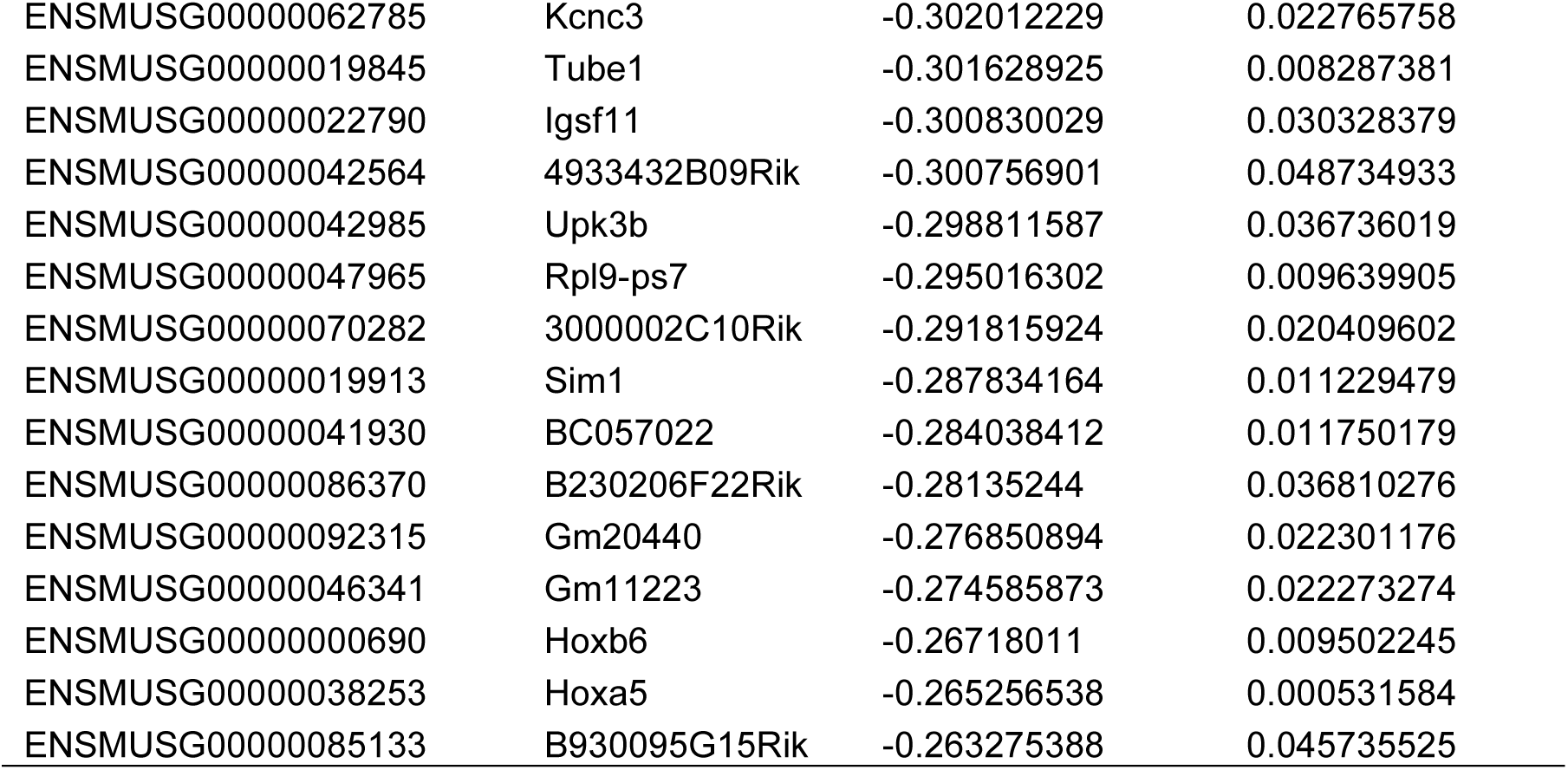
Genes upregulated in *Myf5^Cre/+^;Pbx1^fl/fl;^Pbx2^-/-^* E11.5 embryos (FC<-1.2, P<0.05)

